# CREG1 promotes autophagy and protects the heart against nutritional stress-induced injury and age-associated hypertrophy, fibrosis and diastolic dysfunction

**DOI:** 10.1101/2025.11.11.687936

**Authors:** Yanmei Qi, Russel J. Pepe, Glenn Tejeda, Yechaan Joo, NaYoung K Yang, Kevin B. Hayes, Saum Rahimi, Zhongren Zhou, Shaohua Li, Leonard Y. Lee

**Affiliations:** Departments of Surgery; Department of Pathology and Laboratory Medicine, Rutgers University-Robert Wood Johnson Medical School, New Brunswick, NJ 08901; Rutgers University-New Jersey Medical School, Newark, NJ 07103

## Abstract

**Background:** Cellular repressor of E1A-stimulated genes 1 (CREG1) is an evolutionarily conserved endolysosomal glycoprotein that enhances lysosomal biogenesis and autophagy, suppresses proliferation, and promotes differentiation. A prior gene targeting strategy that produced truncated N-terminal fragments resulted in embryonic lethality, limiting the ability to assess the physiological role of complete CREG1 loss. We hypothesized that CREG1 regulates cardiac autophagy, thereby maintaining cardiac structure and function under both physiological and stress conditions.

**Methods:** We generated true *Creg1* knockout (KO) mice by deleting the entire open reading frame and established a gain-of-function model by inserting human CREG1 into the Rosa26 locus. Cardiac structure and function were assessed in global and cardiomyocyte-specific *Creg1* knockout (cm*Creg1*KO) and knock-in (cm*CREG1*KI) mice. Autophagy was evaluated using biochemical assays, immunofluorescence, electron microscopy, and the CAG-EGFP-RFP-LC3 reporter analysis.

**Results:** Global *Creg1* knockout mice developed progressive cardiac hypertrophy, fibrosis, and diastolic dysfunction at ∼80 weeks of age. At younger ages, CREG1 deficiency increased susceptibility to nutritional stress, resulting in mitochondrial damage and myofiber disruption in cardiomyocytes. cm*Creg1*KO mice exhibited dilated cardiomyopathy, left atrial thrombosis, and lethality around 50 weeks of age; however, interpretation of disease severity is confounded by *Myh6-Cre*–associated cardiotoxicity, which may mask additional pathogenic effects attributable to CREG1 loss. In contrast, cm*CREG1*KI mice demonstrated enhanced exercise capacity under nutritional stress. Mechanistically, CREG1 was localized to endolysosomal and autophagosomal compartments. Loss of CREG1 impaired autophagy flux and mitophagy, likely due to defective autophagosome membrane expansion and degradation. In contrast, CREG1 overexpression enhanced autophagy in cardiomyocytes.

**Conclusions:** CREG1 is a key regulator of cardiac autophagy, protecting the heart against nutritional stress-induced injury and age-associated cardiac hypertrophy, fibrosis, and diastolic dysfunction.

**Graphic Abstract:** 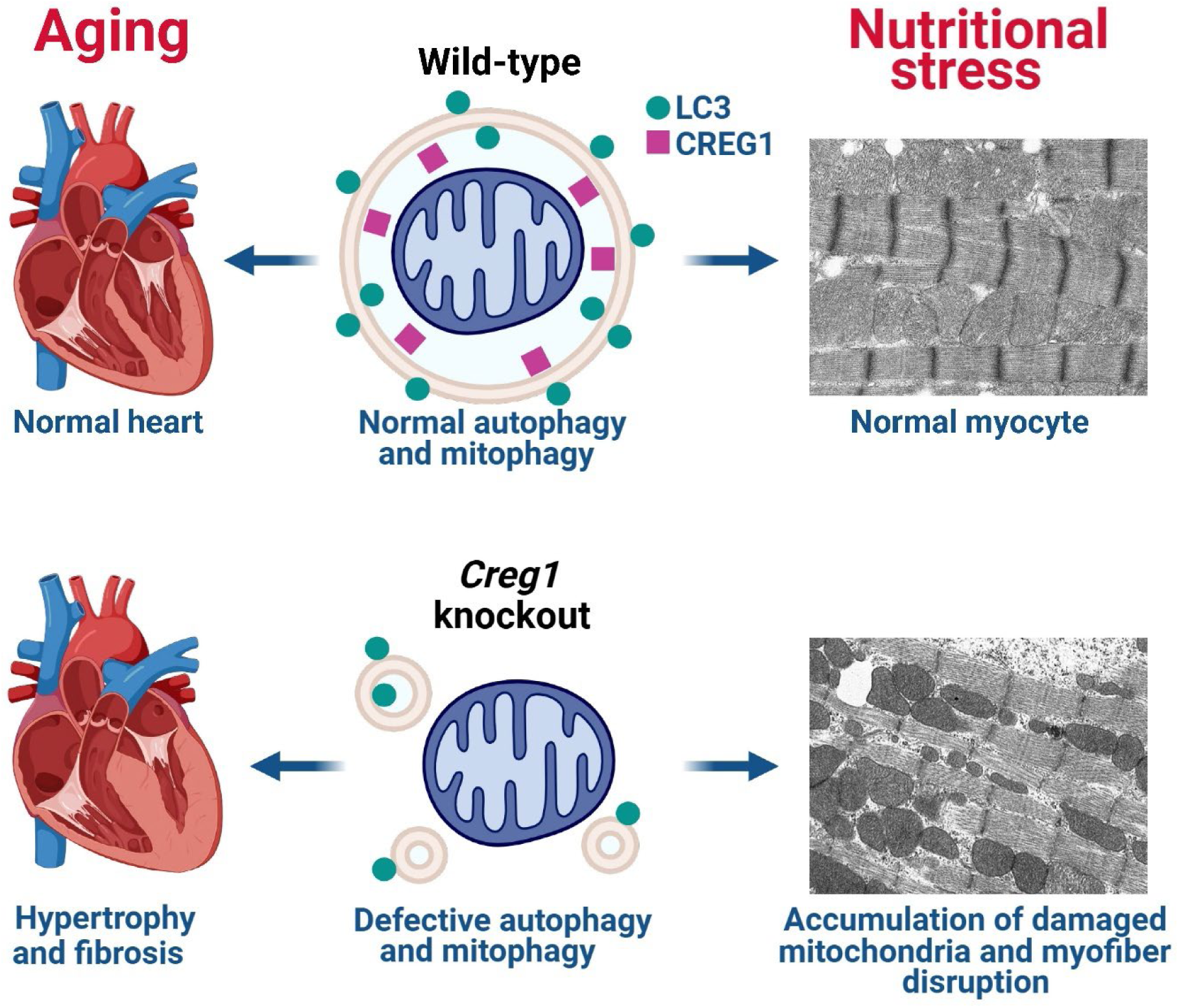

## Introduction

Cellular repressor of E1A-stimulated genes 1 (CREG1) is a small glycoprotein conserved across bacteria, fungi, and animals^1–5^. It contains 220 amino acids in both mice and humans, with two or three N-linked glycosylation sites (N160 and N216 in mice; N160, N193 and N216 in humans) and no identifiable functional domains or motifs^1^. The oligosaccharide chains attached to the glycosylation sites bear mannose-6-phosphate (M6P) that interacted with insulin-like growth factor 2/M6P receptor (IGF2/M6PR) in overlay assay and affinity purification^3, 6, 7^. In cultured cells, CREG1 overexpression inhibited proliferation and transformation while promoting differentiation^1, 3, 8, 9^. Moreover, co-expression of CREG1 with p16^INK4a^ in Li-Fraumeni Syndrome fibroblasts enhanced cellular senescence^10^.

Using knockout/knockdown validated antibodies, we recently demonstrated that CREG1 localizes primarily to endosomes and lysosomes^11^. Our gain- and loss-of-function analyses revealed that CREG1 positively regulates endocytic trafficking, lysosomal biogenesis, and autophagy flux. These findings are consistent with an unbiased protein-protein interaction screen in marine algae, which implicated CREG1 in transferrin-mediated intracellular iron transport^5^. Similarly, a genetic screen in *Drosophila* identified CREG as a regulator of synaptic vesicle size and number, linking it to autism spectrum disorder-associated mutations^12^. Collectively, these studies highlight CREG1 as a critical regulator of endocytic and lysosomal pathways. However, its specific physiological role in mammals remains incompletely understood.

Previous reports also suggest a role for CREG1 in cardiac biology. In a rat model of pressure-overload-induced cardiac hypertrophy, myocardial CREG1 mRNA expression was reduced, whereas expression in kidney, liver, and skeletal muscle remained unchanged^13^. Antisense-mediated suppression of CREG1 in cultured cardiomyocytes enhanced ERK1/2 activation, while cardiomyocyte-specific overexpression of CREG1 driven by the α-myosin heavy chain (α-MHC) promoter in mice inhibited ERK1/2 activation, attenuated cardiac hypertrophy, and improved cardiac function following aortic banding and angiotensin II infusion^14^. Furthermore, *Creg1* heterozygous mice exhibited increased susceptibility to angiotensin-induced myocardial fibrosis at early ages^15^. However, the targeting strategy in this study deleted exons 2 and 3, producing truncated N-terminal fragments (separately reported) and resulting in embryonic lethality^15–17^.

To clarify the role of CREG1 in the heart, we generated true *Creg1* knockout mice by deleting the entire open reading frame. Global deletion caused cardiac hypertrophy, fibrosis, and diastolic dysfunction by 80 weeks of age. Cardiomyocyte-specific deletion under the α-MHC promoter led to dilated cardiomyopathy around 50 weeks. However, interpretation of disease severity is confounded by *Myh6-Cre*–associated cardiotoxicity, which may mask additional pathogenic effects attributable to CREG1 loss. Mechanistically, loss of CREG1 impaired autophagic flux and rendered cardiomyocytes susceptible to injury under nutritional stress, whereas cardiomyocyte-specific CREG1 overexpression enhanced autophagy and improved exercise performance. Together, these findings indicate that CREG1 maintains cardiac structure and function under stress by promoting autophagy.

## Methods

### Animal models

*Creg1^fl/fl^* mice were created using CRISPR-Cas9 by Applied StemCell (Milpitas, CA). Candidate m*Creg1* gRNAs were evaluated *in vitro* by Surveyor assay. Single-stranded oligodeoxynucleotides (ssODNs) targeting the entire open reading frame were inserted into 5’ untranslated region and intron 3-4 in the *Creg1* locus of C57BL/6J mice (forward 5’-TCTCGGAGGGGCGGAACAACGGGCAGGCAATAGTTACCCCGGCAGAGTTTGGTCGGTGGATAACTTCGTATAGCATACATTATACGAAGTTATCGCAGGGACTCCGCAGTCAGCGTCATGGCTGCCCGTGCTCCTGAGCTCGCGCGTTCCCTGC, reverse 5’-TGGCGGCTTTTG TCCAAACGCCCAACCTTGGGAAAAGGTTATTTCTTTGTGGGACCAAGGATAACTTCGTATAGCATACATTATACGAAGTTATAATCTTCAGTCCCGGACCTTAAATTCTTAGTGCTTATGTGTTCTTTGTTTTCATGCTATT). Two independent mouse lines were established, and sequencing confirmed the intended deletion.

*Creg1^fl/fl^* mice were then crossed with *UBC-Cre/ERT2* mice (JAX stock #007001, The Jackson Laboratory)^18^ to generate *Creg1^fl/fl^;UBC-Cre/ERT2* mice. At 8 weeks of age, these mice were injected intraperitoneally with tamoxifen (150 mg/kg for 5 consecutive days) to excise the floxed alleles. *Creg1* knockout mice were subsequently crossed with wild-type mice to produce heterozygotes (*Creg1^+/-^)* lacking the *UBC-Cre/ERT2* transgene, from which homozygous knockouts (*Creg1^-/-^*) were obtained. Cardiomyocyte-specific *Creg1* knockout (cm*Creg1*KO) mice were generated by crossing *Creg1^fl/fl^* mice with *Myh6-Cre* transgenic mice (JAX stock #011038)^19^.

*CREG1* knock-in (KI) mice (*Rosa26^hCREG1^*) were generated by Rutgers University Genome Editing Shared Resource. In the targeting vector, three loxP-flanked stop codons were inserted between the CAG promoter and human *CREG1* cDNA, followed by an IRES and GFP cassette^20^. The vector was inserted into the Rosa26 locus using CRISPR-Cas9 via microinjection into C57BL/6J zygotes. Three independent lines were established and genotyped by PCR. Cardiomyocyte-specific *CREG1* KI (cm*CREG1*KI) mice were generated by crossing *Rosa26^hCREG1/hCREG1^* mice with *Myh6-Cre* transgenic mice.

All animal studies were approved by the Rutgers University Institutional Animal Care and Use Committee. Both male and female mice were used for experiments. Unique IDs were assigned at tagging from a pre-shuffled list. At weaning, same-sex littermates containing both KO/KI and control genotypes were caged together, enrolled sequentially, received identical treatments, and were analyzed concurrently. Runt animals and litters containing only a single genotype were excluded a priori. Daily testing order was randomized independently of genotype. All personnel performing procedures and outcome assessments were blinded to genotype allocation.

### Monitoring autophagy flux using CAG-RFP-EGFP-LC3 reporter mice

Autophagy flux was assessed using *CAG-RFP-EGFP-LC3* dual reporter mice (JAX Stock# 027139, The Jackson Laboratory), in which the LC3 is fused to both red fluorescent protein (RFP) and enhanced green fluorescent protein (EGFP)^21^. *CAG-RFP-EGFP-LC3* mice were crossed with *Creg1^+/-^* mice to obtain *Creg1^-/-^;CAG-RFP-EGFP-LC3* mice and *Creg1^+/+^;CAG-RFP-EGFP-LC3* littermate controls. Mice were fasted for 24-40 hours prior to tissue collection. Hearts were excised, fixed in 4% paraformaldehyde for 1 hour, cryosectioned, and imaged using a Nikon fluorescence microscope.

### Histological analysis

Mouse hearts were fixed in 4% paraformaldehyde, embedded in paraffin, and sectioned at 5 μm thickness. Sections were stained with hematoxylin and counterstained with eosin. Cardiac fibrosis was evaluated by Onestep Trichrome (Green & Red) staining kit (StatLab, McKinney, TX) according to manufacturer’s instructions. Collagen fibers appeared green, nuclei blue, and muscle/cytoplasm red. Images were acquired with a Nikon DS-Fi3 color camera mounted on a Nikon Eclipse TE2000 microscope and processed using NIS-elements imaging software (Nikon).

### Immunofluorescence

Paraffin-embedded mouse heart sections underwent antigen retrieval by boiling in citrate buffer (pH 6.0) for 70 minutes. Primary neonatal mouse cardiomyocytes were fixed with methanol at -20°C for 10 minutes, and immunostaining was performed as previously described ^22^. Images were acquired using a Nikon fluorescence microscope equipped with a Nikon Plan Fluor 40×/1.30 oil objective at 20°C and a Hamamatsu CCD camera. RGB composite images were generated from single-channel captures using NIS-elements software. All images were uniformly sharpened using Unsharp Mask filter in Photoshop (Amount: 200%, Radius: 2.0 pixels).

Cardiomyocyte cross-sectional area was quantified using ImageJ software (NIH, Bethesda, MD). Approximately 150–300 myocytes in left ventricular sections with centrally located nuclei were randomly selected per heart. The cell border highlighted by β-dystroglycan immunofluorescence was detected using ImageJ, and cross-sectional areas were calculated. The mean value per animal was used for statistical analysis.

### Immunoblotting

Mouse tissues and confluent cells were lysed in RIPA buffer. For LC3 and p62 analysis, samples were prepared in SDS lysis buffer containing 1% SDS, 50 mM Tris (pH 7.4), and protease and phosphatase inhibitor cocktails. Immunoblotting was performed as previously described ^23^.

### Electron microscopy

Electron microscopy was performed at Rutgers University-Robert Wood Johnson Medical School Core Imaging Laboratory, as previously described^22^. Heart tissue was fixed in 2.5% glutaraldehyde, 4% paraforamaldehyde and 0.1M sodium cacodylate buffer at 4 °C overnight, followed by washing with 1 mL

0.1 M sodium cacodylate buffer. Samples were then transferred to modified Karnovsky’s fixative for 1 hour and postfixed in 1% osmium tetroxide for 1 hour. After a graded series of ethanol dehydration and Epon/SPURR resin infiltration and embedding, thin sections of resin-embedded tissues were cut with a diamond knife in an ultramicrotome, stained, and examined with a Philips CM12 electron microscope. Images were captured with an AMT-XR11 digital camera.

### Echocardiography

Transthoracic echocardiography was performed using a high-resolution Vevo 3100 ultrasound system (VisualSonics, Toronto, Canada) equipped with a 30 MHz linear transducer at Rutgers Molecular Imaging Center. Mice were anesthetized with 1.5–2% isoflurane in 100% oxygen and placed on a temperature-controlled imaging platform to maintain body temperature at 37 °C. Hair was removed from the chest with a depilatory cream, and ECG electrodes were attached to the limbs for continuous heart rate monitoring. Two-dimensional guided M-mode images were obtained in the parasternal short-axis view at the level of the papillary muscles. Left ventricular (LV) end-diastolic and end-systolic diameters (LVEDD and LVESD) were measured, and fractional shortening (FS) was calculated as (LVEDD–LVESD)/LVEDD × 100%. LV ejection fraction (EF) was determined using standard algorithms provided by the software. All measurements were averaged from at least three consecutive cardiac cycles and analyzed in a blinded manner.

### Blood pressure monitoring

Blood pressure was monitored using a CODA tail-cuff system (Kent Scientific, Torrington, CT). Mice were placed in restraint tubes on a pre-warmed platform for 15 minutes prior to measurement. An occlusion cuff was positioned at the base of the tail, and a volume-pressure recording sensor cuff was placed adjacent to it. The occlusion cuff was inflated to 250 mmHg and then slowly deflated over 20 seconds. Each measurement session consisted of 15-25 inflation/deflation cycles, with the first 5 cycles designated as acclimation and excluded from analysis. Data from the remaining cycles were averaged to calculate systolic, diastolic, and mean arterial pressures. To improve reproducibility and reduce stress-induced variability, mice are acclimated to the procedure for three consecutive days before data collection.

### Data-independent acquisition (DIA) mass spectrometry

Proteomic analysis was performed at Rutgers University Biological Mass Spectrometry Facility using DIA mass spectrometry as previously described^24^. Peptide samples were separated by reverse-phase liquid chromatography and analyzed on a high-resolution mass spectrometer operated in DIA mode. Raw DIA data were processed with Skyline-daily software using a spectral library. Protein identification and quantification were performed with a false discovery rate threshold of 1% at both peptide and protein levels.

### Mouse neonatal cardiomyocytes

Mouse neonatal cardiomyocytes were isolated from 1-day-old mice via trypsin digestion and cultured as previously described^25, 26^. Cells were serum-starved for 24 hours before fixation for immunofluorescence microscopy.

### Treadmill exercise endurance

An acute exhaustive exercise test was conducted using a six-lane rodent treadmill (Ugo Basile, Comerio, VA, Italy). After a three-day acclimation period, mice ran on a level treadmill (0° incline) starting at 10 m/min for 3 minutes. The speed was then increased by 4 m/min every 3 minutes until exhaustion. Exhaustion was defined as the inability to resume running after remaining on the low-intensity shock grid (1.5 mA, 200 ms pulses, 4 Hz) for five consecutive seconds. Time to exhaustion and total running distance were recorded.

## Results

### Global deletion of *Creg1* leads to age-dependent cardiac hypertrophy, myocardial fibrosis, and diastolic dysfunction

To elucidate the physiological function of CREG1, we generated *Creg1* floxed mice using CRISPR technology by inserting loxP sites to flank exons 1 to 3 that encode the entire open reading frame (separately reported). To delete *Creg1* in adults, we crossed *Creg1^fl/fl^*mice with *UBC-Cre/ERT2* mice to obtain *Creg1^fl/fl^ ;UBC-Cre/ERT2* mice, which were injected intraperitoneally with tamoxifen (150 mg/kg for 5 consecutive days) at 8 weeks of age to induce excision of the floxed alleles. We next generated heterozygous mice (*Creg1^+/-^*) by crossing the *Creg1* knockout mice with wild-type C57BL/6 mice. Intercrosses between the *Creg1^+/-^* mice lacking the *Cre* transgene yielded *Creg1^-/-^* offspring at Mendelian ratios, suggesting that CREG1 is dispensable for development (Figs. 1A, 1B and data reported separately). This result is different from the previous reports by targeting exons 2 and 3, which led to embryonic lethality^15–17^.

**Figure 1.**
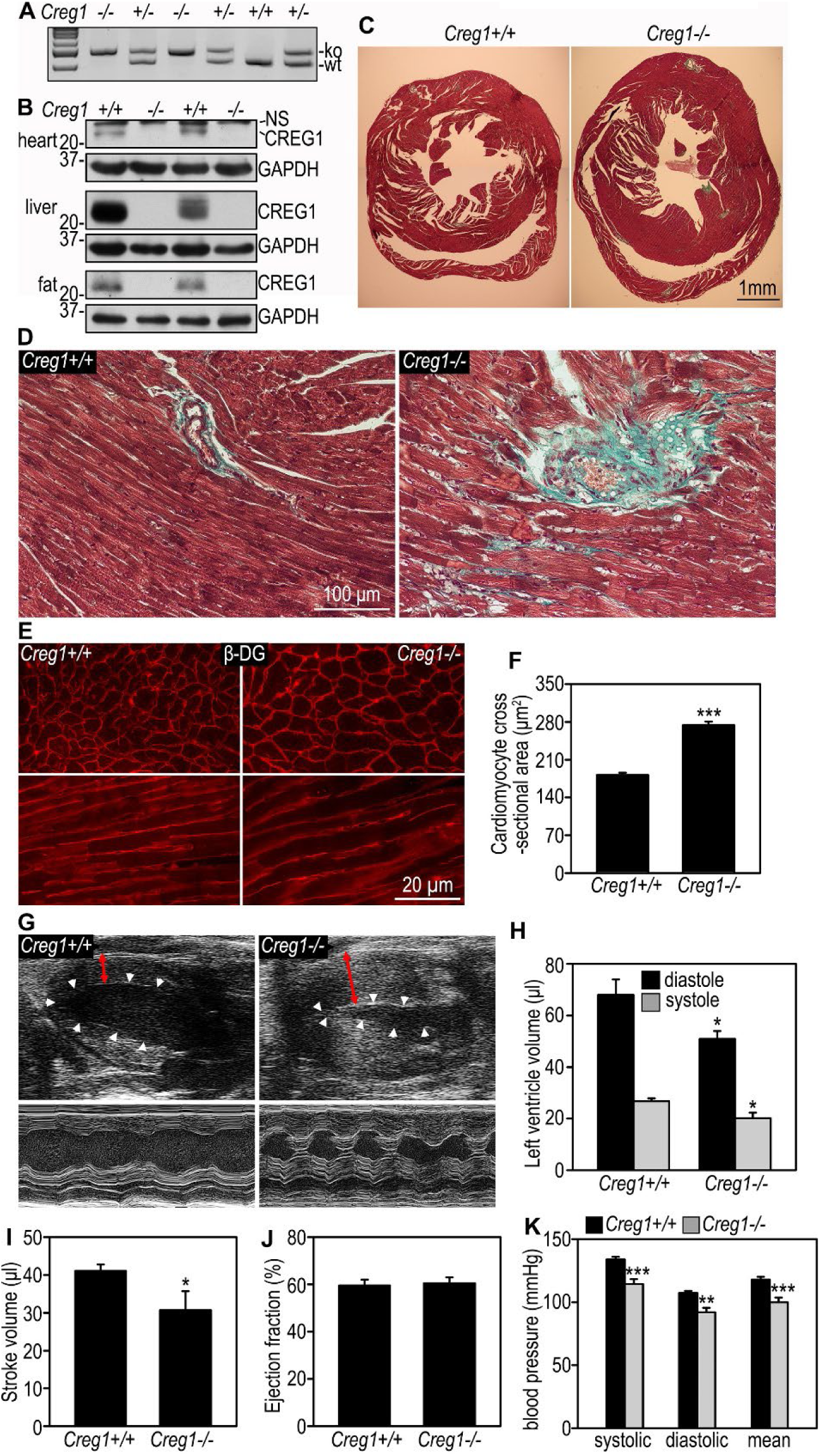
Global deletion of *Creg1* leads to age-dependent cardiac hypertrophy, myocardial fibrosis, and diastolic dysfunction. **(A)** Representative PCR genotyping of *Creg1* wild type (wt, +/+), knockout (ko, -/-) and heterozygotes (+/-) mice. **(B)** Immunoblots show the absence of the CREG1 protein in the heart, liver, and fat of *Creg1*KO mice. GAPDH served as a loading control. **(C)** Cross-section of the heart from a 123-week-old *Creg1*KO mouse shows concentric hypertrophy compared to its littermate control. **(D)** Trichrome staining in (C) indicates increased perivascular and interstitial fibrosis (green) in the *Creg1*KO heart. **(E)** β-Dystroglycan (β-DG) immunostaining of heart cross-sections (upper) and longitudinal sections (lower) highlighted the plasma membrane and revealed increased cardiomyocyte diameter in *Creg1*KO mice compared to controls. **(F)** Cardiomyocyte cross-sectional areas were significantly increased in *Creg1*KO mice compared to littermate controls (***P < 0.001 Student’s *t*-test; *Creg1*^+/+^: n = 4, *Creg1*^-/-^: n = 5). **(G)** Representative B-mode (upper) and M-mode (lower) echocardiographic images of a 103-week-old *Creg1*KO mouse and its littermate control reveal a thickened left ventricular wall and left ventricular outflow tract obstruction during systole in *Creg1*KO mice. Arrowheads indicate the endocardium, while red arrows mark anterior wall thickness at the end of diastole. **(H, I and J)** Quantification of echocardiographic measurements showed significantly reduced end-diastolic left ventricular volume (H) and stroke volume (I) in *Creg1*KO mice compared to littermate controls, with no significant change in ejection fraction (J) (*P < 0.05 Paired *t*-test; n = 4 litters, 4 *Creg1*^+/+^ and 6 *Creg1*^-/-^ mice). (**K)** *Creg1^-/-^* mice displayed significantly lower systolic, diastolic, and mean blood pressures compared with their littermate controls (n = 4 litters; **P<0.01, ***P<0.001; paired *t*-test).

*Creg1^-/-^* mice did not show any discernible phenotypic differences compared to littermate controls. However, necropsy revealed that the hearts of knockout mice older than 80 weeks were enlarged, with significantly thickened left ventricles (Fig. 1C). Trichrome staining on heart tissue sections showed mild fibrotic tissue deposition around arteries and veins in control mice, with cardiomyocytes aligned in parallel in longitudinal sections (Fig. 1D). In contrast, *Creg1^-/-^* hearts displayed markedly increased fibrosis, not only in perivascular areas but also between cardiomyocytes. In addition, myofiber alignment was disrupted, especially near fibrotic regions surrounding the vessels. Of note, cardiomyocytes in knockout hearts were significantly enlarged. To quantify the cross-sectional area of cardiomyocytes, we immunostained for β-dystroglycan (β-DG), a basement membrane receptor highly expressed on the surface of cardiomyocytes^27^. As shown in Figs 1E and 1F, the cross-sectional area of cardiomyocytes increased by 50% in knockout hearts. Collectively, these results suggest that CREG1 ablation leads to cardiac hypertrophy and myocardial fibrosis in aged mice.

Next, we performed a functional analysis of *Creg1* knockout hearts. B-mode and M-mode echocardiography revealed left ventricular wall thickening, reduced diastolic chamber volume, and occasional left ventricular outflow tract obstruction in *Creg1* knockout mice (Fig. 1G). Statistical analysis demonstrated significantly decreased diastolic and systolic left ventricle volume and stroke volume in knockout hearts (Figs. 1H and 1I), while the EF remained unchanged (Fig. 1J). These findings indicate that *Creg1* knockout hearts develop diastolic dysfunction with preserved ejection fraction.

Hypertension is the most common cause of left ventricular hypertrophy^28^. To determine whether CREG1 ablation affects blood pressure, we monitored 37-week-old mice for 3 weeks using the CODA non-invasive rodent blood pressure monitor system. Both systolic and diastolic pressures were significantly lower in *Creg1^-/-^* mice compared to their *Creg1^+/+^*littermates, suggesting the left ventricular hypertrophy observed in *Creg1*-deficient hearts is not due to elevated blood pressure.

### *Creg1* knockout mice are susceptible to nutritional stress-induced cardiac injury

Autophagy is upregulated in the heart and is essential for cardiomyocyte survival and the maintenance of cardiac function under stress conditions such as starvation and myocardial ischemia^29^. We previously showed that CREG1 depletion in hepatocytes reduces lysosomal enzyme levels, impairs lysosomal acidification, and disrupts autophagic flux^11^. To investigate the role of CREG1 in cardiac adaptation to nutritional stress, 35-week-old mice were subjected to a 40-hour fast, a standard method to induce autophagy. After fasting, *Creg1^-/-^* mice were markedly less active than their *Creg1^+/+^* littermates. Histological analysis of *Creg1^-/-^* heart revealed cardiomyocyte vacuolation, degeneration, and reduced eosin staining (Figure 2A). Ultrastructural examinations showed that mitochondria in *Creg1^+/+^*cardiomyocytes were regularly aligned between the myofibrils and associated with small lipid droplets (Figure 2B). In contrast, mitochondria in *Creg1^-/-^* cardiomyocytes appeared condensed, irregular, and disorganized, and were frequently associated with enlarged lipid droplets. Sarcomere length increased by 38%, and myofibrillar disruption was evident in knockout hearts (Figures 2B and 2C). Heart-to-body ratios were also examined in adult mice after an overnight fast. Under ad libitum conditions, ratios were slightly lower in *Creg1^-/-^*mice (Figure 2D). Fasting modestly decreased heart-to-body ratios in *Creg1^+/+^* mice, likely reflecting autophagy activation, whereas *Creg1^-/-^* mice displayed a significant increase. Together, these findings suggest that CREG1 deficiency sensitizes the heart to nutritional stress-induced injury.

**Figure 2.**
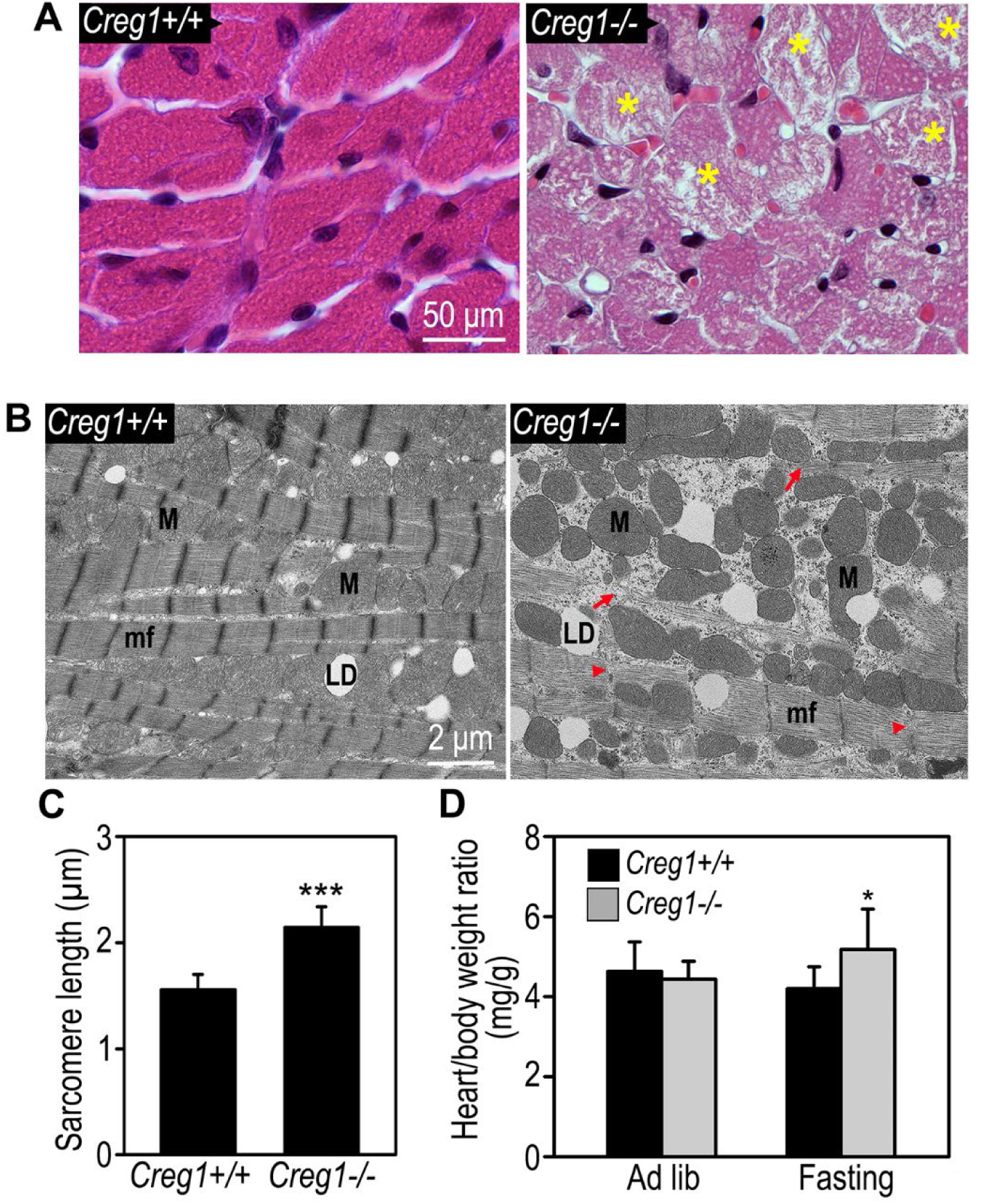
*Creg1* knockout mice are susceptible to nutritional stress-induced cardiac injury. **(A)** Representative H&E staining of heart cross-sections from *Creg1*^-/-^ mice and littermate control *(Creg1*^+/+^) following overnight fasting. Yellow * indicates cardiomyocyte vacuolation. **(B)** Electron micrographs of hearts from a 4-month-old *Creg1*^-/-^ mouse and its littermate control (*Creg1*^+/+^) following overnight fasting. Red arrows denote disrupted myofibers; red arrowheads indicate damaged Z-discs in *Creg1*^-/-^cardiomyocytes. LD: lipid droplet; M: mitochondrion; mf: myofiber. **(C)** Sarcomere length was significantly increased in *Creg1^-/-^* cardiomyocytes (n = 66 sarcomeres for each group; ***P<0.001; Student’s *t-*test). **(D)** After an overnight fast, heart-to-body weight ratios were significantly increased in *Creg1^-/-^*mice compared with littermate controls (n = 9 litters, *P<0.05; paired *t-*test).

### *Myh6-Cre*-mediated cardiomyocyte-specific deletion of *Creg1*

To further investigate the role of CREG1 in the heart, we generated cm*Creg1*KO mice by crossing *Creg1^fl/fl^* mice with *Myh6-Cre* transgenic mice^19^. In cm*Creg1*KO hearts, CREG1 protein levels were reduced by 82.4±10.8% (mean ± SD, N = 4 pairs) compared to *Creg1^fl/fl^* littermate controls (Figure 3A). In sharp contrast to *Creg1^fl/fl^* mice, which survived throughout the observation period, all cm*Creg1*KO mice died of heart failure with respiratory distress between 45 and 55 weeks of age (Figure 3B). Necropsy revealed significantly dilated heart chambers and increased heart-to-body weight ratios (Figures 3C and 3D). Left atria were largely occupied by what appeared to be organized thrombi.

**Figure 3.**
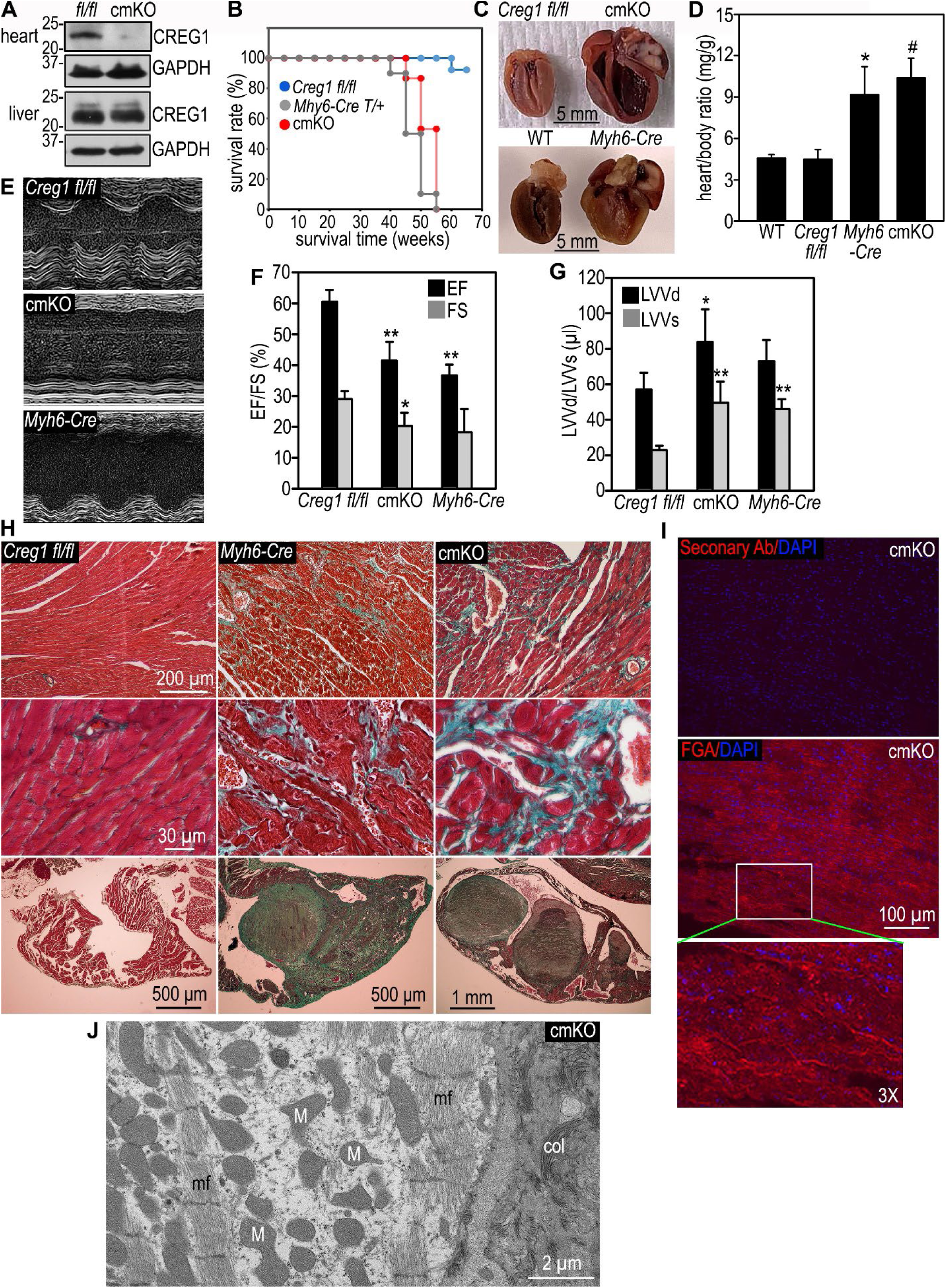
Analysis of cardiomyocyte-specific *Creg1* knockout (cm*Creg1*KO) mice. **(A)** Immunoblots show substantial reduction of CREG1 expression in the heart, but not in the liver, of cm*Creg1*KO mice. **(B)** Kaplan-Meier survival curves show premature lethality of both cm*Creg1*KO and Myh*6-Cre* mice at 40-55 weeks, whereas most *Creg1^fl/fl^* control mice survived over the observed period. **(C)** Representative cross-sections of the hearts showing all four chambers in *Myh6-Cre*, cm*Creg1*KO and the control mice. Both *Myh6-Cre* and cm*Creg1*KO displayed dilated left and right ventricles and left atrial thromboses, with cm*Creg1*KO mice showed more severe dilation and thinner interventricular septum than *Myh6-Cre mice*. **(D)** The heart/body weight ratio was increased in both *Myh6-Cre* (\**P* < 0.05) and cm*Creg1*KO (^#^*P* < 0.05) mice compared to wild-type (WT) and *Creg1^fl/fl^* controls. The cm*Creg1*KO mice showed a slightly higher ratio than the Myh*6-Cre* mice. **(E, F and G)** Echocardiograms show decreased left ventricle contraction, reduced ejection fraction (EF), and increased left ventricular volumes at the end of diastole (LVVd) and systole (LVVs) in both *Myh6-Cre* and cm*Creg1*KO mice (*P<0.05, **P<0.01 compared with *Creg1^fl/fl^* mice). **(H)** Trichrome staining showed myocardial fibrosis (green) and chronic left atrial thrombosis in both cm*Creg1*KO and *Myh6-Cre* mice. cm*Creg1*KO hearts appeared more severe. **(I)** Immunostaining for fibrinogen A (FGA) confirmed left atrial thrombosis in cm*Creg1*KO mice. The lower panel is a 3× magnification of the boxed area. **(J)** Representative electron micrograph of cm*Creg1*KO hearts following overnight fasting. M: mitochondrion; mf: myofiber; col: collagen.

Echocardiography performed at 42-43 weeks on cm*Creg1*KO mice showed marked left ventricular dilation and systolic dysfunction (Figures 3E-3G). Both EF and FS were significantly reduced, while left ventricular end-diastolic and systolic volumes were increased. Trichrome staining revealed severe myocardial fibrosis and disorganization of ventricular myofibers, with the left atrium extensively infiltrated by collagen fibers stained in green (Figure 3H). Immunostaining for the fibrinogen α chain confirmed thrombosis in the left atrium (Figure 3I). At the ultrastructural level, we observed condensed and irregular mitochondria, disrupted myofibrils, and collagen fiber accumulation (Figure 3J). These findings suggest that cm*Creg1*KO mice develop dilated cardiomyopathy, left atrial thrombosis, and ultimately heart failure.

Because expression of the *Myh6-Cre* transgene has been associated with myocardial damage^30, 31^, we included age-matched wild-type and *Myh6-Cre* mice as additional controls. Wild-type mice showed survival rates and heart morphology comparable to those of *Creg1^fl/fl^* mice. However, *Myh6-Cre* mice died around the same age as cm*Creg1*KO mice, though ventricular dilation, myocardial fibrosis, and left atrial thrombosis appeared less severe. These results suggest that *Myh-Cre*-mediated cardiomyocyte-specific gene deletion may not be a suitable approach for studying age-dependent cardiac abnormalities in CREG1-deficient mice.

### CREG1 ablation impairs cardiomyocyte autophagy

To assess the effect of nutritional stress on cardiac CREG1 expression, C57BL/6J mice were fasted for 16 hours. Immunoblot analysis revealed a 2.2-fold increase in cardiac CREG1 expression after fasting (Figure 4A). Because nutritional stress activates autophagy to support cardiomyocyte survival under these conditions^32^, we next examined whether loss of CREG1 affects autophagy by immunostaining for LC3B, an autophagosome-associated protein. Numerous LC3B puncta were readily detectable in wild-type hearts after a 40-hour fast (Figure 4B), but they were significantly reduced in *Creg1^-/-^* hearts (Figures 4B and 4C). Autophagic flux analysis confirmed the reduction of LC3B-II, which is associated with the LC3B puncta, in the absence of CREG1 (Figure 4D). Subsequent injection of the lysosomal inhibitor chloroquine to block autophagy flux resulted in sustained lower levels of LC3B-II in *Creg1^-/-^*hearts, along with an increase in p62, a marker for lysosomal degradation of ubiquitinated cargoes. Parallel findings were noted in cm*Creg1*KO hearts (Figure 4E). These results suggest that the diminished LC3B-II levels and LC3B puncta in the absence of CREG1 are caused by reduced autophagosomal formation rather than increased degradation.

**Figure 4.**
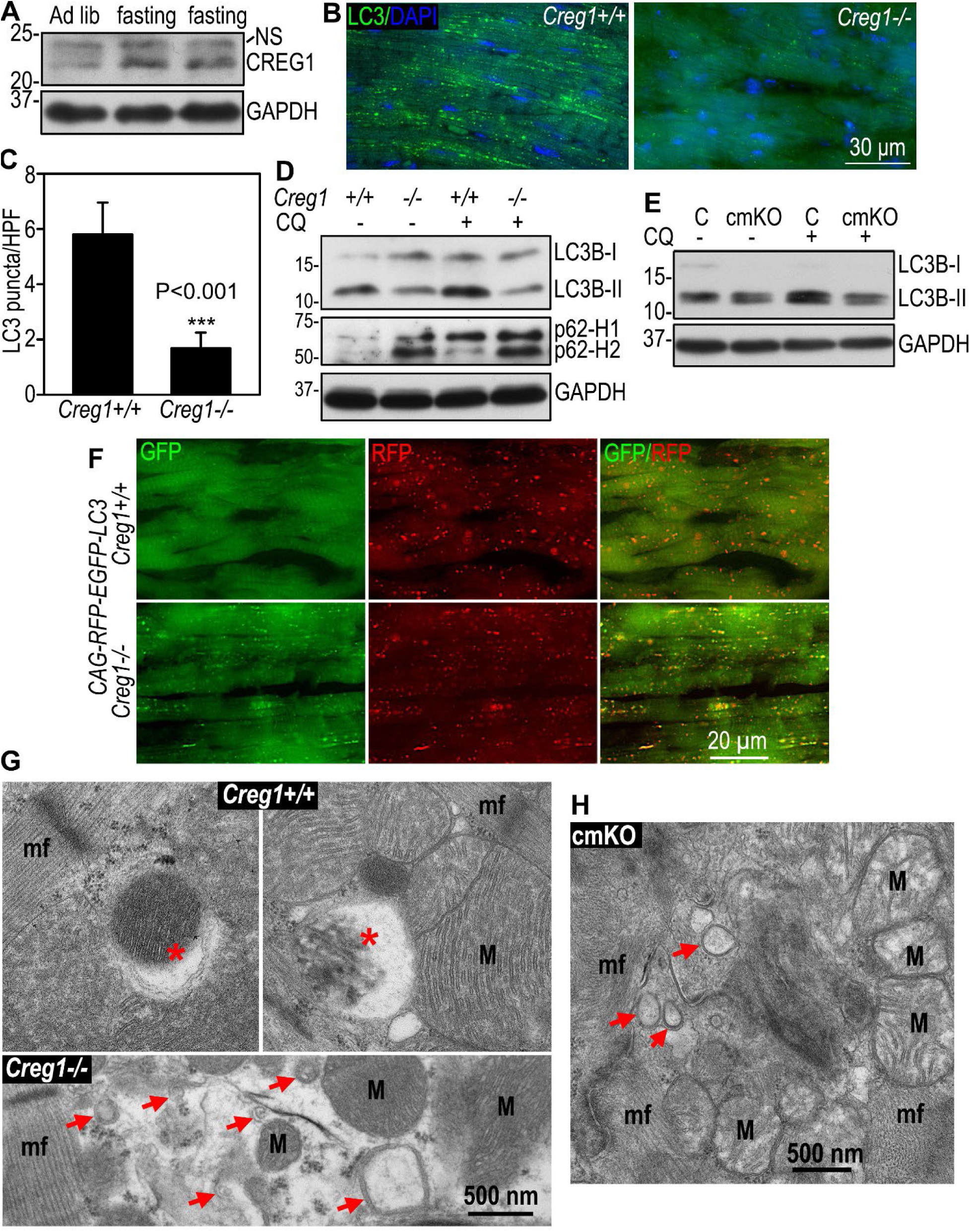
CREG1 ablation impairs autophagy flux and mitophagy in cardiomyocytes. **(A)** CREG1 expression was increased in the heart after 16-hour fasting. **(B)** LC3 immunofluorescent puncta were significantly reduced in *Creg1*^-/-^ hearts after 24-hour fasting. **(C)** Quantification of LC3 puncta per high-power field (HPF) using a 40× objective. *Creg1^+/+^*vs *Creg1^-/-^*, student’s *t*-test ***P<0.001. **(D)** Immunoblots show decreased LC3BII/I ratios and increased p62 levels in *Creg1*^-/-^ hearts with or without chloroquine (CQ) treatment compared to littermate controls. **(E)** Immunoblot analysis demonstrated decreased LC3BII/I ratio in cm*Creg1*KO heart compared to littermate controls. **(F)** *Creg1*^+/+^ and *Creg1*^-/-^mice were crossed with *CAG-RFP-EGFP-LC3* reporter mice and fasted for 40 hours. *Creg1*^-/-^ heart tissues exhibited predominantly yellow fluorescence puncta, suggesting impaired autophagosome degradation, while in littermate controls, most LC3 puncta appeared red, indicating efficient autolysosomal formation. **(G)** Electron microscopy showed autophagosomes containing damaged mitochondria (red asterisks) in *Creg1*^+/+^ cardiomyocytes. Red arrows indicate mini autophagosomes in *Creg1*^-/-^ cardiomyocytes. **(H)** Mini autophagosomes were also observed adjacent to damaged mitochondria in cm*Creg1*KO cardiomyocytes.

Although chloroquine administration led to LC3B-II accumulation in control hearts (Figures 4D and 4E), it did not increase LC3B-II levels in *Creg1* knockout hearts. This suggests that autophagosomal degradation is also compromised. To further investigate autophagy flux in *Creg1^-/-^* hearts, we crossed *Creg1^-/-^* mice with *CAG-RFP-EGFP-LC3* transgenic mice. This system utilizes the stability of RFP in acidic pH, contrasting with the quenching of EGFP in acidic lysosomes. This distinction allows for the identification of autophagosomes (neutral pH) and autolysosomes (acidic pH)^21, 33, 34^. After a 40-hour fast, the majority of LC3B puncta appeared red in wild-type hearts, indicating autolysosome formation (Figure 4F). In contrast, approximately half of the LC3B puncta appeared yellow in *Creg1^-/-^* hearts, further indicating impaired autophagosomal degradation. Notably, the introduction of RFP-EGFP-LC3 in the knockout hearts led to an increase in LC3B puncta (Figures 4B and 4F), suggesting a partial rescue of autophagosome formation.

Adult cardiomyocytes are long-lived, post-mitotic cells that are abundant in mitochondria due to their high energy demand for contraction. Consequently, they are uniquely vulnerable to the accumulation of malfunctioning mitochondria^35, 36^. Mitophagy plays a central role in maintaining mitochondrial homeostasis and reducing cellular oxidative stress^37–39^. We conducted electron microscopy on heart tissues to assess the role of CREG1 in mitophagy. In wild-type mice subjected to a 40-hour fast, mitophagy in cardiomyocytes was readily discernible (Figure 4G). In contrast, *Creg1^-/-^* and cm*Creg1*KO hearts exhibited only small, closed double-membrane vesicles or cup-shaped membranes, ranging in size from 100 to 300 nm, adjacent to damaged mitochondria (Figures 4G and 4H). This observation suggests potential deficiencies in autophagosomal membrane expansion in the absence of CREG1.

### CREG1 is localized to endolysosomal compartments, and its ablation downregulates proteins associated with endocytic trafficking and autophagy

To elucidate the molecular mechanisms by which CREG1 regulates autophagy, we employed next generation DIA mass spectrometry. We identified a total of 6,184 proteins, with 288 (110 upregulated and 178 downregulated) exhibiting significant changes in the heart of 17∼18-week-old *Creg1^-/-^* mice compared to wild-type littermates following 24-hour fasting (P<0.05, Figure 5A). DAVID gene functional classification analysis revealed the largest group of proteins involved in intracellular trafficking (Figure 5B). Among the downregulated proteins, we identified ATG9A and Osbpl2, known participants in lipid transfer and autophagosomal membrane expansion. The functional association of CREG1 with endolysosomal compartments was further supported by immunofluorescence analysis which showed partial co-localization of CREG1 with the early endosomal marker Rab5, the late endosomal marker Rab7, and the lysosomal marker LAMP-1 in primary cardiomyocytes isolated from neonatal hearts (Figure 5C). CREG1 was also detected in the endoplasmic reticulum marked by calreticulin. Notably, when cardiomyocytes were serum-starved for 24 hours, CREG1 partially co-localized with LC3B puncta (Figure 5D). These findings suggest that CREG1 may be involved in autophagosome membrane expansion by regulating endocytic trafficking.

**Figure 5.**
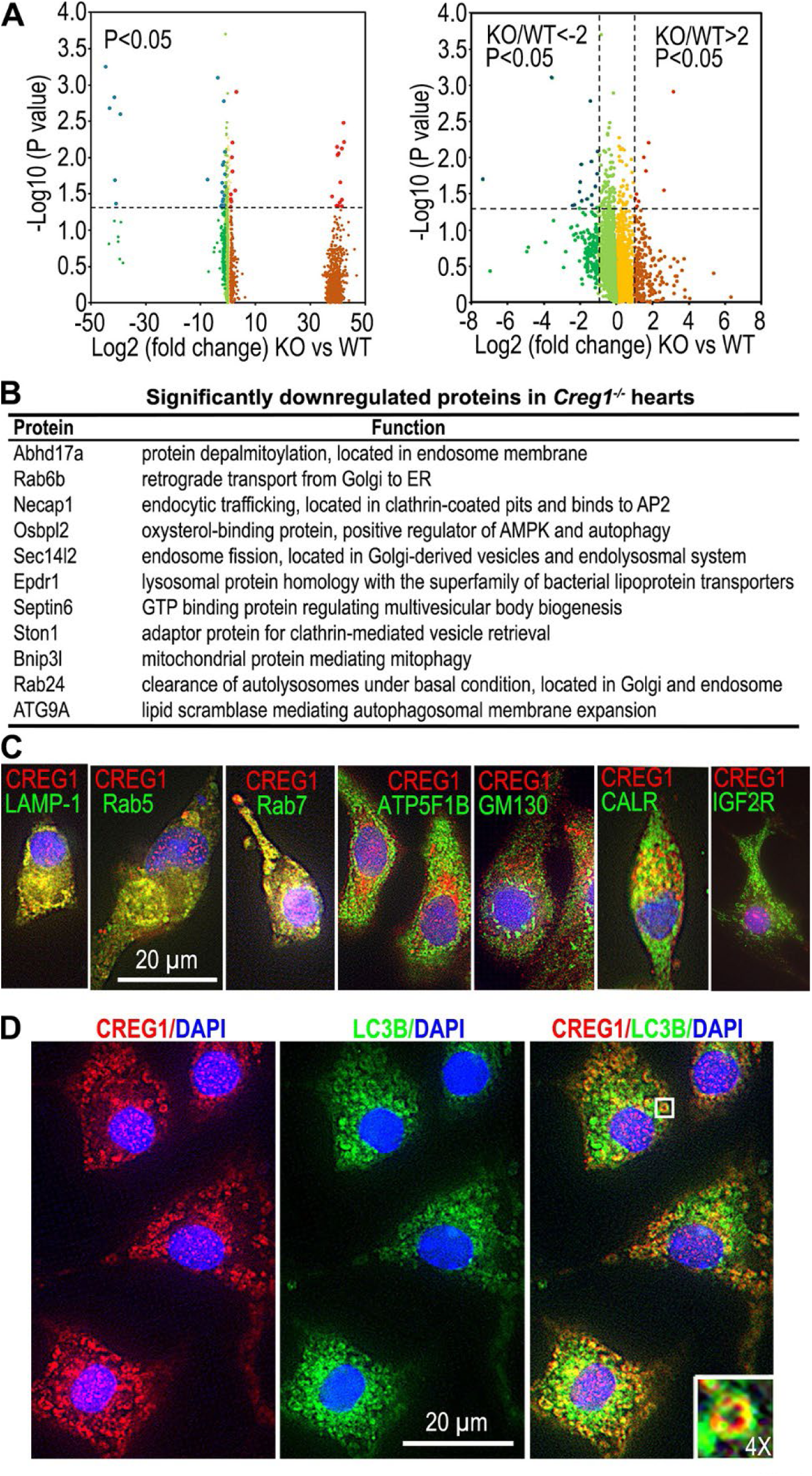
CREG1 is associated with autophagosomes in cardiomyocytes and CREG1 ablation downregulates proteins involved in endocytic trafficking, lysosomal function and autophagosomal membrane expansion. **(A)** Volcano plots comparing the proteomic profile of heart lysates from *Creg1*^-/-^(KO) and control mice (WT). Each point represents a single protein; significantly upregulated and downregulated proteins are highlighted in red and blue, respectively. N = 4 littermate pairs. **(B)** A list of significantly downregulated proteins in the hearts of *Creg1*^-/-^ mice compared to wild-type (WT) controls. These proteins were identified by quantitative proteomic analysis and met the cutoff criteria of log2 KO/WT <-2, and adjusted P < 0.05. **(C)** Co-immunostaining for CREG1 and markers for lysosomes (LAMP-1), early endosomes (Rab5), late endosomes (Rab7), mitochondria (ATP5F1B), the Golgi apparatus (GM130), the endoplasmic reticulum (CALR), and IGF2R in primary mouse cardiomyocytes. **(D)** Immunofluorescence microscopy showed partial co-localization of CREG1 with the autophagosome marker LC3B in primary mouse cardiomyocytes after 24-hour serum deprivation. The inset is a 4× magnification of the boxed area.

### Cardiomyocyte-specific overexpression of CREG1 improves exercise endurance and enhances autophagy flux

To investigate whether cardiomyocyte-specific overexpression of CREG1 enhances autophagy and improves stress endurance, we generated conditional *CREG1* KI mice by inserting human *CREG1* cDNA into the Rosa26 locus. cm*CREG1*KI was achieved by crossing *Rosa26^hCREG1/hCREG1^* mice with *Myh6-Cre* transgenic mice to remove the stop codons inserted between the CAG promoter and CREG1 cDNA (Figures 6A and 6B). Immunoblotting showed a 3.1-fold increase in CREG1 expression in the heart of heterozygous cm*CREG1*KI mice compared to wild-type and *Myh6-Cre* mice (Figure 6C). The cm*CREG1*KI mice were born healthy and fertile and were indistinguishable from control *Rosa26^hCREG1^* littermates. HE staining revealed no abnormalities in the KI heart at 15 weeks of age (data not shown). Compared to wild-type mice, cm*CREG1*KI mice were more active under fasting conditions. *Myh6-Cre* mice exhibited significantly reduced run time and distance in a maximal treadmill exercise test compared with wild-type littermates (Figures 6D and 6E). In contrast, after a 16-hour fast, cmCREG*1*KI mice, despite carrying the detrimental *Myh6-Cre* transgene, ran significantly longer and farther than *Rosa26^hCREG1^* littermates, which lacked the *Myh6-Cre* transgene and performed similarly to wild-type mice. These findings suggest that the overexpression of CREG1 in cardiomyocytes improves exercise endurance under stress, likely by enhancing cardiac function.

**Figure 6.**
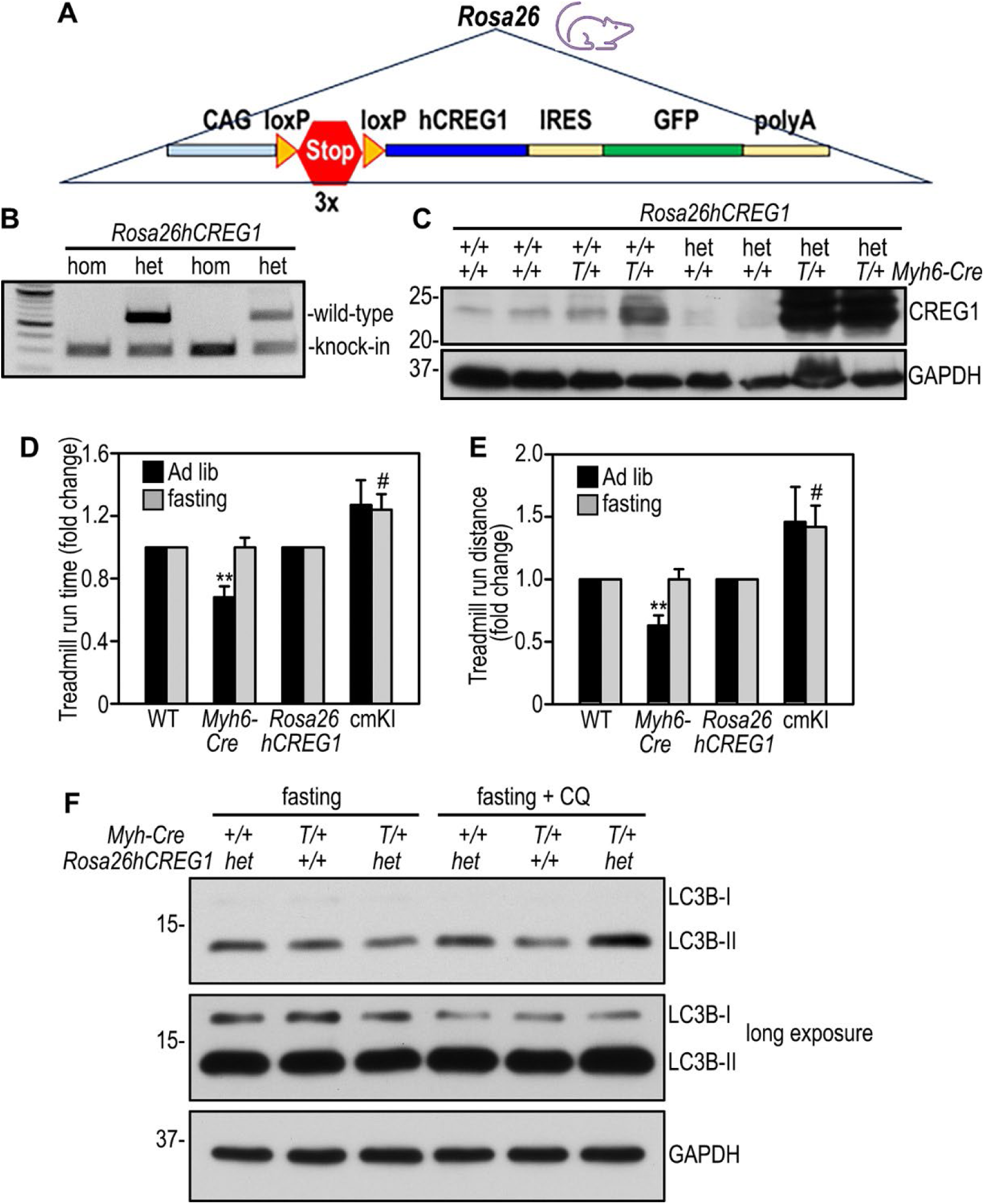
Cardiomyocyte-specific CREG1 knock-in improves treadmill performance under nutritional stress and enhances cardiac autophagy flux. (A) **The CREG1 knock-in strategy for conditional expression of human *CREG1* cDNA in C57BL/6J mice. (B) Representative PCR genotyping of homozygous (hom) and heterozygous (het) *CREG1* knock-in mice. (C) Immunoblot analysis confirmed CREG1 overexpression in the heart of cm*Creg1*KI mice. GAPDH served as a loading control. (D and E) Treadmill endurance test revealed a significant decrease in running time and distance in *Myh6-Cre* mice with Ad libitum feeding, while cm*CREG1*KI (cmKI) mice exhibited markedly enhanced endurance following overnight fasting, compared to their respective litter controls (mean ± SE; student’s *t*-test, *Myh6-Cre* vs WT **P < 0.01; cmKI vs *Rosa26^hCREG1^* ^#^P < 0.05). (F) Immunoblots show increased LC3BII/I ratios in cm*CREG1*KI hearts after a 40-hour fast, compared to their littermate controls.**

Next, we examine the effects of cardiomyocyte-specific CREG1 overexpression on autophagy flux. Mice were fasted for 40 hours followed by intraperitoneal administration of chloroquine to inhibit lysosomal activity. Immunoblotting showed that LC3B-II levels were lower in cmCREG*1*KI than *Rosa26^hCREG1^* and *Myh6-Cre* control hearts in the absence of chloroquine (Figure 6F). Following chloroquine administration, LC3B-II levels in cmCREG*1*KI hearts were markedly higher than those of control hearts, likely due to increased production and accumulation in lysosomes and autophagosomes after lysosomal inhibition. These findings suggest that CREG1 overexpression enhances both autophagosomal formation and degradation, which may contribute to improved exercise performance under nutritional stress.

## Discussion

The present study demonstrates that CREG1 promotes cardiac autophagy and protects the heart against nutritional stress-induced injury. Several lines of evidence support this conclusion: (1) CREG1 expression was significantly upregulated in the heart during starvation; (2) global and cardiomyocyte-specific deletion of *Creg1* inhibited LC3 lipidation in the heart, both with and without lysosomal inhibition by chloroquine, and led to p62 accumulation, indicating impaired autophagosome formation; (3) CAG-RFP-EGFP-LC3 reporter analysis revealed yellow puncta in CREG1-deficient hearts, consistent with a blockade in autophagosome-lysosome fusion and/or lysosomal dysfunction; (4) electron microscopy showed mini, empty autophagosomes adjacent to damaged mitochondria with impaired mitophagy in knockout hearts; (5) CREG1-deficient cardiomyocytes were more susceptible to starvation-induced injury; and (6) Cardiomyocyte-specific overexpression of CREG1 enhanced autophagic flux and improved exercise performance under nutritional stress.

Autophagy is a conserved catabolic pathway involving lysosome-mediated degradation of misfolded proteins and damaged organelles, thereby maintaining cardiac structure and function under normal and stress conditions^40–44^. Disruption of autophagy in the heart leads to cardiomyopathy, as evidenced by cardiomyocyte-specific deletion of *Atg5*, which results in dilated cardiomyopathy and heart failure^45–49^. Likewise, mutations in the lysosome-associated membrane protein-2 (LAMP-2) gene cause Danon disease, a disorder characterized by severe dilated or hypertrophic cardiomyopathy^50, 51^. LAMP-2 deficiency in mice induces autophagic vacuole accumulation, cardiac hypertrophy, and contractile dysfunction of cardiac muscles at late stages^52^. Our global *Creg1* knockout mice displayed a similar but milder phenotype, suggesting that impaired autophagy contributes to the observed pathology.

The mechanisms by which CREG1 regulates cardiac autophagy remain incompletely understood. Our autophagic flux analyses and CAG-RFP-EGFP-LC3 imaging suggest that both autophagosome formation and degradation are compromised in the absence of CREG1. The presence of mini autophagosomes near damaged mitochondria in CREG1-deficient cardiomyocytes indicates a defect in autophagosomal membrane expansion. A similar phenotype was reported in Hela cells harboring mutations in ATG9A, the only transmembrane protein in the core autophagy machinery, which functions as phospholipid scramblase to facilitate autophagosomal membrane growth via ATG9A vesicles^53–55^. Consistent with this, we observed partial co-localization of CREG1 with markers for the endoplasmic reticulum, early and later endosomes, and autophagosomes. Moreover, mass spectrometry revealed downregulation of Osbpl2, a lipid transfer protein^56^, in CREG1-deficient hearts. These findings raise the possibility that CREG1 regulates autophagosome biogenesis through ATG9A- and/or Osbpl2-dependent pathways, a hypothesis warranting further investigation.

Previous studies suggested that CREG1 interacts with IGF2R, a multifunctional receptor involved in IGF2 clearance and M6P-dependent trafficking of lysosomal proteins^57^. While CREG1 is expected to enhance IGF2R internalization and thereby accelerate IGF2 degradation^58, 59^, our findings argue against IGF2R as a major mediator of CREG1 function in the heart. Although we detected co-immunoprecipitation of CREG1 with IGF2R in adult mouse hearts (data not shown), the two proteins did not co-localize in primary cardiomyocytes (Figure 5C). Moreover, phenotypic differences between *Creg1* and *Igf2r* knockout mice underscore their functional divergence: *Igf2r* knockout mice are perinatal lethal, approximately 30% larger than controls, and exhibited marked cardiac overgrowth due to elevated IGF2^60, 61^. In contrast, *Creg1* knockout mice are viable, slightly smaller in early life (data not shown), and display normal heart-to-body ratios (Figure S1A). Consistently, proliferating cell nuclear antigen (PCNA) expression was not significantly altered in CREG1-deficient neonatal hearts (Figure S2A and S2B). Collectively, these data suggest that IGF2R is unlikely to mediate the cardioprotective role of CREG1.

Cardiac aging is characterized by hypertrophy, fibrosis, and the accumulation of damaged mitochondria, leading to diastolic dysfunction^62–64^. Impaired autophagy plays a critical role in this process, whereas enhancing autophagy and mitophagy has been shown to confer cardioprotection and extend the lifespan in mice^47^. The impaired autophagy and mitophagy observed in *Creg1* knockout mice likely exacerbates age-related cardiac hypertrophy and fibrosis. Of note, this phenotype is unlikely to result from pressure overload, as blood pressure was reduced compared with littermate controls (Figure 1K). We cannot exclude the possibility that systemic metabolic disturbance, such as hyperglycemia and insulin resistance observed in global *Creg1* knockout mice (reported separately), also contributes to the cardiac phenotype. Another limitation is the known cardiotoxicity of the *Myh6-Cre* transgene, which has been reported previously and further characterized in our study^31, 65–67^, restricting analysis of cardiac structure and function beyond 45 weeks of age. Despite these constraints, two key findings strongly support the cardioprotective role of CREG1 under nutritional stress: (1) young CREG1-deficient mice were more susceptible to starvation-induced cardiac injury, and (2) cm*CREG1*KI mice exhibited enhanced exercise performance.

It is noteworthy that a recent study reported muscle-specific deletion of *Creg1* in mice using the muscle creatine kinase promoter, which appeared to enhance autophagy and mitophagy, although autophagy flux was not directly assessed^68^. The authors showed massive mitochondrial vacuolization in skeletal muscle after CREG1 loss, resembling changes induced by chemical compounds, bacterial and viral toxins, or osmotic stress. However, because the gene-targeting approach in that study generated truncated N-terminal fragments and resulted in embryonic lethality, the observed increase in mitophagy may have been secondary to mitochondrial damage caused by these fragments. This highlights the critical importance of careful gene-targeting design to ensure reliable models.

## Acknowledgments

The authors thank Peter Romanienko and Ghassan Yehia for generating the Rosa26*^hCREG1^* mice, Rajesh Patel for electron microscopy services, and Haiyan Zheng for mass spectrometry services. Gene functional classification analysis was performed using the DAVID Gene Functional Classification Tool.

## Abbreviations

CREG1: cellular repressor of E1A-stimulated genes 1; M6P: mannose-6-phosphate; IGF2/M6PR: insulin-like growth factor 2/M6P receptor; α-MHC: α-myosin heavy chain; ssODNs: single-stranded oligodeoxynucleotides; cm*Creg1*KO: cardiomyocyte-specific *Creg1* knockout; cm*CREG1*KI: cardiomyocyte-specific *CREG1* knock-in; EGFP: enhanced green fluorescent protein; RFP: red fluorescent protein; LC3: microtubule-associated protein 1A/1B-light chain 3; LVEDD: left ventricular end-diastolic diameter; LVESD: left ventricular end-systolic diameter; FS: fractional shortening; EF: ejection fraction; DIA: data-independent acquisition; β-DG: β-dystroglycan; LAMP-2: lysosome-associated membrane protein-2; PCNA: proliferating cell nuclear antigen

## Novelty and Significance

### What Is Known?

**•** CREG1 is small glycoprotein localized in the endolysosomal compartment and involved in endocytic trafficking and lysosomal biogenesis.
**•** CREG1 overexpression in cultured cells suppresses proliferation and promotes differentiation and senescence.

### What New Information Does This Article Contribute?

**•** *Creg1* knockout mice develop age-associated cardiac hypertrophy, fibrosis, and diastolic dysfunction.
**•** CREG1 deficiency increases cardiac susceptibility to nutritional stress-induced injury.
**•** Loss of CREG1 disrupts cardiac autophagy flux and mitophagy, likely through impaired autophagosomal membrane expansion and autophagosome degradation.
**•** Cardiomyocyte-specific overexpression of CREG1 enhances autophagy flux and improves exercise endurance during nutritional stress.

A prior *Creg1* gene targeting strategy resulted in embryonic lethality due to production of truncated N-terminal fragments, leaving the physiological role of CREG1 unresolved. Here, we generated a complete *Creg1* knockout and a Rosa26 knock-in model overexpressing human CREG1 to investigate its cardiac function in vivo. Integrated gain- and loss-of-function analyses revealed that CREG1 is essential for maintaining autophagy-mediated mitochondrial homeostasis, preserving myocardial structure, and sustaining cardiac performance under both normal and stress conditions. These findings establish CREG1 as a critical regulator of cardiac health and stress resilience.

---

**Figure S1.**
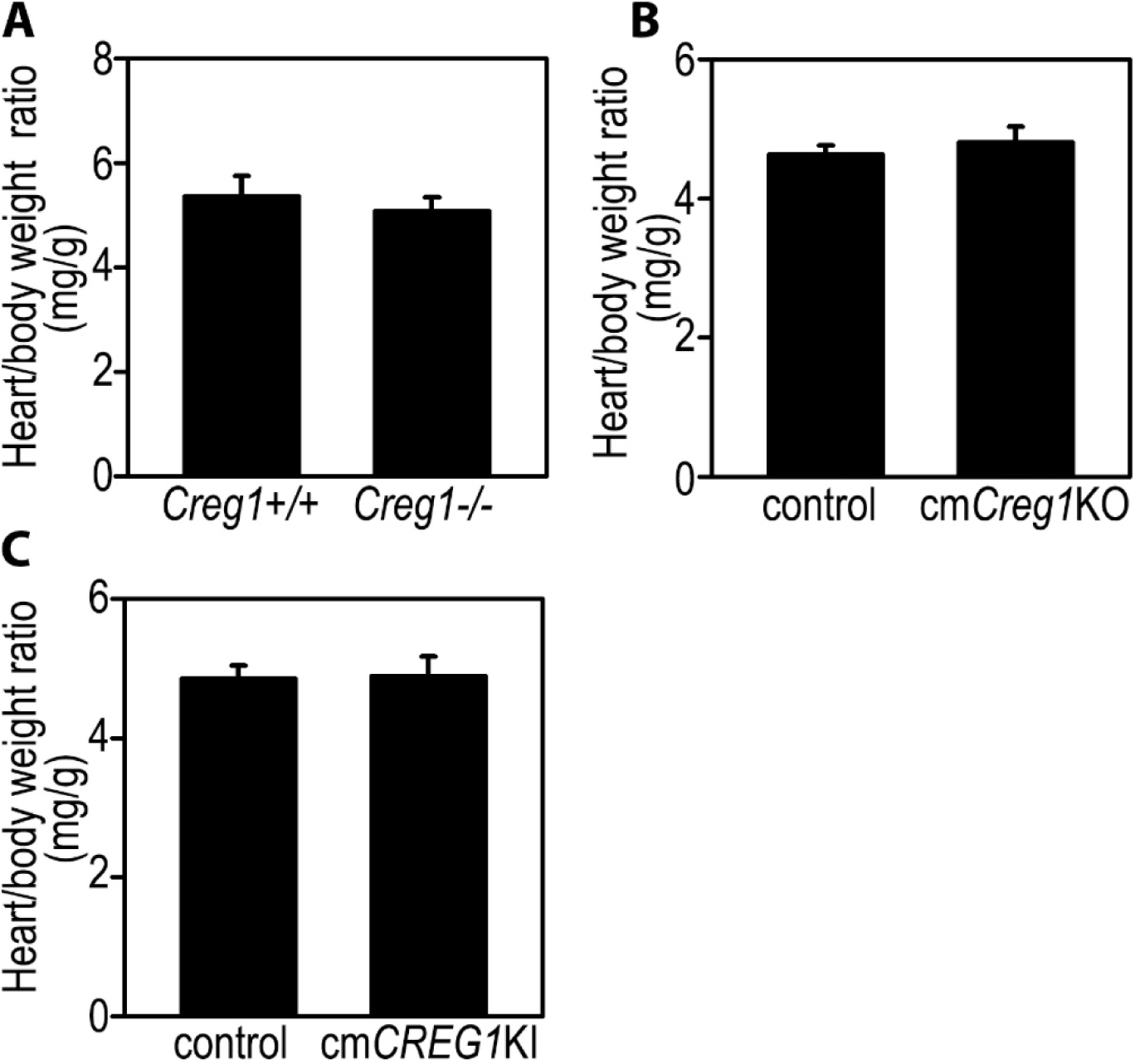
Heart-to-body weight ratios. Heart-to-body weight ratios of *Creg1^-/-^*, cm*Creg1*KO, and cm*CREG1*KI mice were not significantly altered compared with littermate controls.

**Figure S2.**
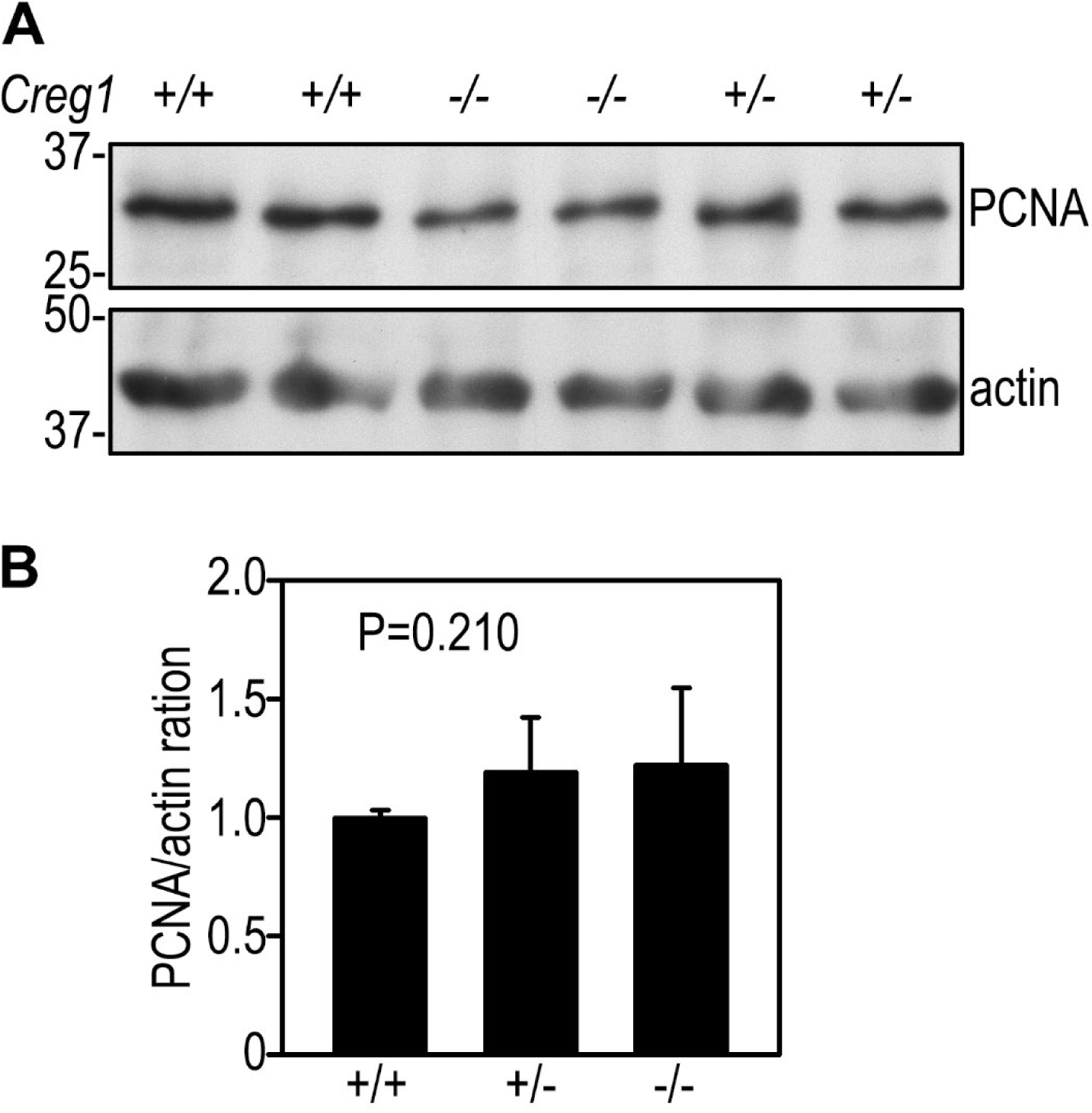
Proliferating cell nuclear antigen (PCNA) expression. Cardiac PCNA was analyzed in P0.5 neonates by immunoblotting and was not significantly changed in *Creg1^-/-^* mice compared with *Creg1^+/+^* and *Creg1^+/-^* littermate controls (N = 4 for each group).

